# Healthy adipocytes provide a protective environment by limiting viral infection through cell-cell communication

**DOI:** 10.1101/2025.09.11.675622

**Authors:** Alexsia Richards, Max Friesen, Troy W. Whitfield, Lee Gehrke, Rudolf Jaenisch

## Abstract

Adipocytes, long regarded primarily as energy storage cells, are increasingly recognized as active players in immune regulation. In metabolic disorders such as obesity and diabetes—conditions characterized by adipocyte dysfunction—patients often exhibit worsened outcomes following viral infections. However, the role of healthy adipocytes in modulating antiviral immunity remains poorly understood. Here, we demonstrate that healthy adipocytes secrete factors that confer antiviral protection to neighboring cells. We developed a fully human pluripotent stem cell (hPSC)-derived system including adipocytes, immune, and vascular cells to evaluate the antiviral capacity and infectivity of various cell types. Our cell system has the advantage of being of human origin, as opposed to animal models. Through our technological development, we have removed cell culture medium as a variable by adapting all cell types to a single base medium. We found that only adipocytes could induce an antiviral state in adjacent mural and immune cells. This unique immunomodulatory capacity is mediated, at least in part, by the STING-dependent secretion of low levels of interferon-alpha (IFN-α) from healthy adipocytes. Notably, pharmacological induction of metabolic dysfunction in adipocytes diminished their antiviral activity, revealing a previously unrecognized link between metabolic health and antiviral defense. These findings identify a novel role for adipocytes in orchestrating local antiviral responses and provide new insight into how metabolic dysfunction may compromise host defense during viral infections.

## Introduction

Flaviviruses, including West Nile virus (WNV), Zika virus (ZV), and Ilheus virus (IV), are emerging global health threats, with infections on the rise due to factors such as expanding mosquito populations, urbanization, and climate change^1^. These viruses present significant public health challenges, as they can cause severe diseases ranging from fever to encephalitis and meningitis^2^. Flavivirus infection begins with the bite of an infected tick or mosquito, introducing the virus into the skin and bloodstream. These viruses initially infect local immune cells at the bite site, such as dendritic cells and macrophages. These infected immune cells then facilitate the spread of the virus to the lymphatic system and bloodstream, leading to systemic dissemination^3^. Once in circulation, flaviviruses have the potential to infect multiple cell types, including the endothelial cells lining blood vessels as well as perivascular cells like pericytes and smooth muscle cells ^4–8^.

Adipocytes are increasingly recognized for their immune-modulating functions, beyond their traditional role in energy storage^9^. Notably, perivascular adipose tissue, a specialized form of adipose tissue that surrounds blood vessels, acts as a critical signaling hub by releasing bioactive molecules, including cytokines and adipokines, which can modulate the behavior of nearby vascular and immune cells^10, 11^. Metabolically unhealthy conditions, such as obesity and lipodystrophy, are often associated with worse outcomes during viral infections, emphasizing the importance of adipose tissue in shaping immune responses^12, 13^. While the exact mechanisms for this phenomenon are not exactly clear, it is likely mediated by both a general deterioration in health in metabolically unhealthy individuals, as well as a loss of beneficial and protective properties of healthy adipose tissue. Given this well-established role in impacting adjacent cells, we hypothesized that metabolically healthy adipocytes actively contribute to the response of vascular and immune cells to viral infection.

Although murine models offer valuable insights into adipocyte biology, significant metabolic and immunological differences exist between mouse and human fat cells, limiting the direct translation of findings. Additionally, the pathology observed in mouse models of viral infection often does not correlate with what is observed in human patients^14, 15^. Conversely, the use of primary human adipocytes is often hindered by limited availability, donor variability, and challenging culture conditions, all of which affect experimental reproducibility. To overcome these challenges, we established a system utilizing exclusively human pluripotent stem cell (hPSC)-derived cells, including adipocytes, smooth muscle cells (SMCs), pericytes (PCs), macrophages (MACs), dendritic cells (DCs), and endothelial cells (ECs). Through culture condition optimizations, we were able to culture all these cell types in a single, shared medium, thereby removing this variable from all the viral experiments. We exposed hPSC-derived target cells to adipocyte secreted factors before viral infection. Exposure to adipocyte-secreted factors significantly reduced infection in both vascular and immune cells, highlighting a broad protective role. We further show that this antiviral activity is mediated through adipocyte secretion of IFNα. Notably, even without external stimuli, adipocytes maintain high baseline levels of interferon regulatory factor (IRF) activation, creating an antiviral environment in neighboring cells. By demonstrating the protective role of healthy adipocytes during viral infections, this study underscores the importance of metabolic health in shaping antiviral immune responses.

## Results

### Adipocytes secrete factors that reduce infection in vascular and immune cells

To investigate the potential antiviral effects of adipocyte-secreted factors, we generated a panel of hPSC-derived target cells, including SMCs, PCs, ECs, MACs, and DCs, based on their susceptibility to infection by all three flaviviruses (**Fig. S1**). We then exposed these target cells to adipocyte-conditioned medium for 24 hours and subsequently infected the cells with a panel of flaviviruses, including West Nile virus (WNV), Zika virus (ZV), and Ilheus virus (IV), and quantified the viral replication **(Fig. 1A)**. This approach allowed us to isolate the impact of adipocyte-secreted factors on viral infection of target cells without the confounding effects of adipocyte infection. Our results show that pretreatment of hPSC-derived SMCs with adipocyte-conditioned medium significantly reduced infectious virus production, whereas exposure to control medium conditioned by adipocyte precursors had no effect (**Fig. 1B, C)**. To determine if adipocytes were unique in their antiviral activity, we pretreated hPSC-derived SMCs with either adipocyte-conditioned medium or medium conditioned by hPSC-derived ECs, hPSC-derived PCs, or primary human fibroblasts. We observed that only adipocyte-conditioned medium had an antiviral effect **(Figure 1D)**. The antiviral impact of adipocyte conditioned medium was not limited to SMC infection, as pre-treatment with adipocyte conditioned medium also significantly reduced subsequent infection by WNV, IV, and ZV of hPSC-derived pericytes **(Figure 1E-G)**. To determine if a decrease in the number of infected cells was responsible for the reduced production of infectious virus in our target cells, we quantified the number of infected SMCs following treatment with either unconditioned medium, progenitor-conditioned medium, or adipocyte- conditioned medium. Treatment with adipocyte-conditioned medium significantly reduced the number of infected cells, suggesting that exposure to adipocyte-secreted factors decreases the overall infection rate (**Fig. 1H, I**). However, we cannot exclude the possibility that the amount of infectious virus released from each infected cell is also reduced.

**Figure 1.**
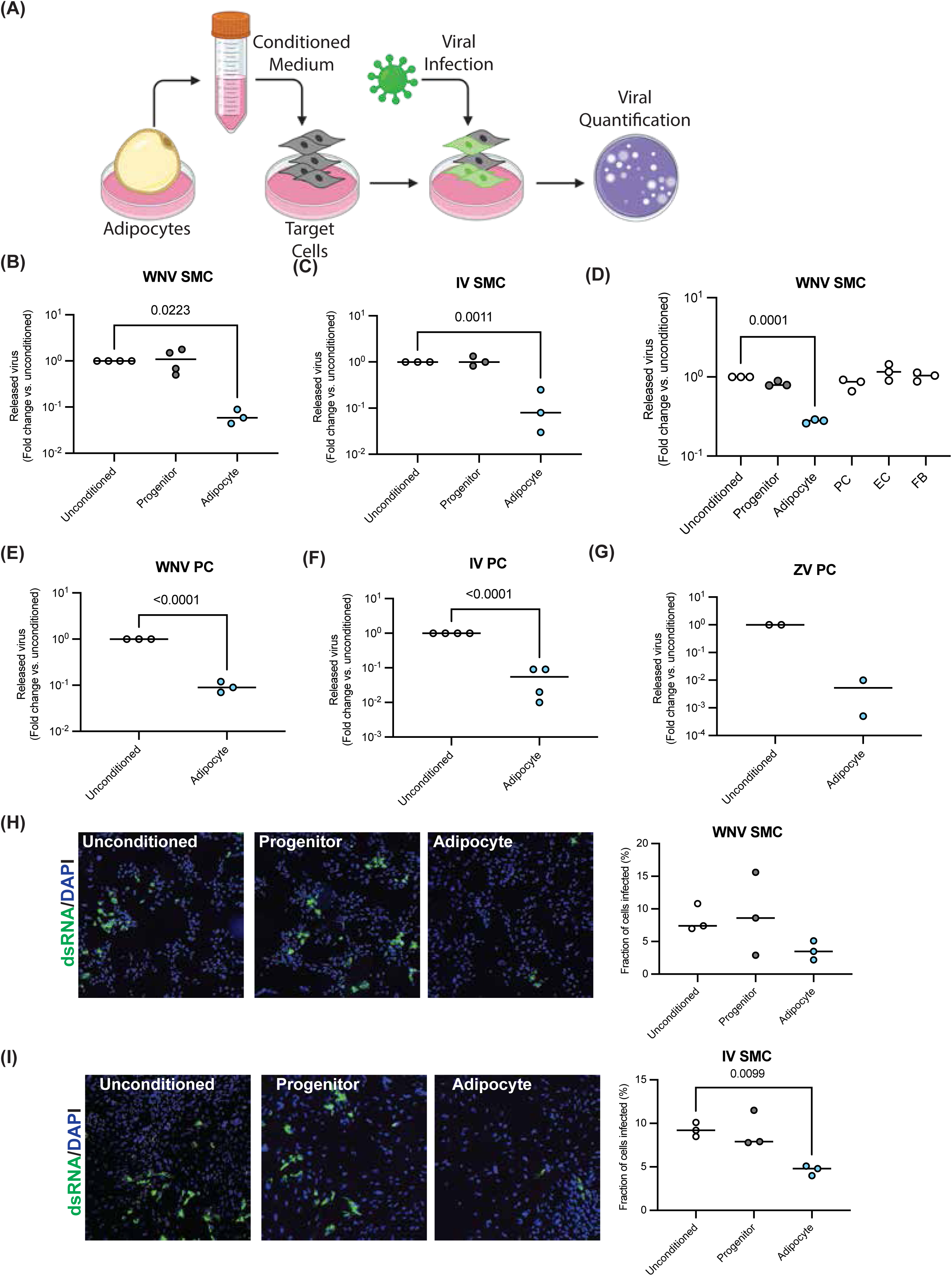
Adipocytes secrete factors that restrict infection of mural c ells: **(A)** Schematic description of the experimental setup. **(B,C)** Plaque assays showing released virus from hPSC-derived smooth muscle cells treated with unconditioned, progenitor-conditioned or adipocyte-conditioned medium prior to infection with West Nile Virus (B) or Ilheus Virus (C). **(D)** Plaque assay quantitation of released virus from smooth muscle cells treated with unconditioned, progenitor-, adipocyte-, pericyte-, endothelial cell-, or fibroblast-conditioned medium prior to infection with West Nile virus. **(E-G)** Plaque assays showing released virus from hPSC-derived pericytes treated with unconditioned or adipocyte-conditioned medium prior to infection with West Nile Virus (E), Ilheus Virus (F), or Zika Virus (G). **(H,I)** Immunofluorescence staining of smooth muscle cells treated with unconditioned, progenitor-conditioned or adipocyte- conditioned medium prior to infection with West Nile Virus (H) or Ilheus Virus (I). Cells were fixed and stained with an antibody against double-stranded RNA (dsRNA) at 48 hours post infection. For all experiments three experimental replicates were performed. Infectious virus release was quantitated at 48 hours post-infection. Mean values for each experimental replicate were normalized to the mean value of unconditioned or progenitor medium condition for the same experimental replicate.

Immune cells are critical target cells during viral infection and have the potential to spread viral infection systemically^16^. To determine whether exposure to adipocyte- secreted factors could reduce the infection of immune cells, we similarly exposed hPSC-derived macrophages and dendritic cells to either adipocyte-conditioned medium or medium conditioned by adipocyte progenitor cells prior to infection. Our results show that exposure to adipocyte-conditioned media significantly reduced subsequent infection compared to treatment with progenitor-conditioned media **(Figure S2A-C)**. Similar to our observations in hPSC-derived SMCs, treatment with conditioned media from hPSC-derived PCs, ECs, and primary fibroblasts had no impact on subsequent viral infection **(Figure S2D)**. We expanded our studies to hPSC-derived DCs and observed that treatment with adipocyte-conditioned media significantly reduced subsequent infection with WNV relative to treatment with progenitor-conditioned media **(Figure S2E)**.

To assess whether the observed antiviral effect extended beyond flaviviruses, we repeated the experiment using SARS-CoV-2. hPSC-derived human liver organoids (HLOs) and hPSC-derived SMCs have both been shown to be susceptible to SARS-CoV-2 infection^17, 18^. Pretreatment of hPSC-derived SMCs with adipocyte-conditioned medium significantly reduced the release of infectious SARS-CoV-2 as well as the number of infected cells. In contrast, treatment with progenitor conditioned media had no effect **(Fig. S3A-B).** This data suggests that the antiviral activity of adipocyte- secreted factors is not limited to flaviviruses.

Interestingly, we found that not every cell type benefited from the protective effect of adipocyte-conditioned medium. Perivascular and immune cells showed a significant reduction in viral infection after adipocyte conditioned medium treatment, however, ECs, brain microvascular endothelial cells (BMECs), and HLOs showed no benefit (**Fig. S4).** We speculate that the lack of protective effect in these cell types is that either they are not receptive to the protective factors in the adipocyte-conditioned medium, or they already produce those factors themselves.

To ensure that the observed reduction in infectivity was not the result of reduced target cell viability, we assessed the cell viability of hPSC-derived SMCs, PC, MACs, and DCs following 72 hours of culture in either unconditioned medium, adipocyte-conditioned medium, or each cell type’s standard culture medium. Our results show that culturing of target cells in adipocyte-conditioned medium did not reduce cell viability **(Fig. S5)**.

To assess whether direct exposure to adipocytes would recapitulate the antiviral effects we observed following treatment with adipocyte conditioned medium, we established a co-culture system using Transwells, where hPSC-derived PCs, MACs, or DCs shared medium with hPSC-derived adipocytes **(Fig. 2A)**. After co-culture for 24 hours, the target cells were infected apically with WNV **(Fig. 2A)**. Similar to our observations with adipocyte conditioned medium, the presence of adipocytes reduced WNV production in PCs, MACs, and DCs **(Fig. 2B-D)**. Notably, the adipocytes themselves appeared resistant to viral infection, as evidenced by the low amount of infectious virus released from the adipocyte-only condition compared to PCs or MACs **(Fig. 2C-D)**.

**Figure 2.**
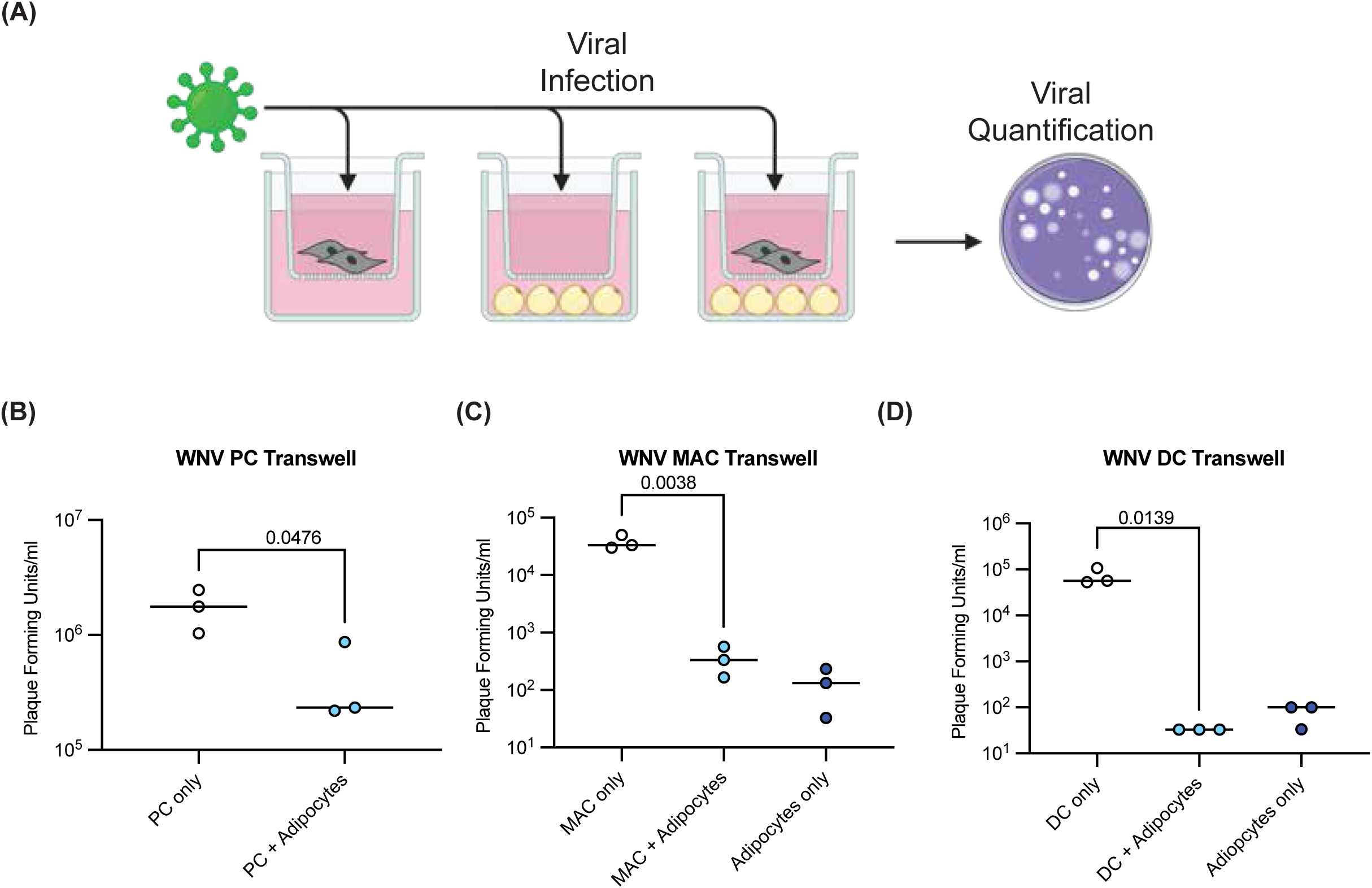
Paracrine signaling from adipocytes reduces infection of neighboring cells: **(A)** Schematic description of the experimental setup. **(B-D)** Adipocytes were plated in the bottom of the transwell and either pericytes (PCs) (B), macrophages (MACs) (C) or dendritic cells (DCs) (D) were plated in the top chamber. 24 hours after the initiation of co-culture West Nile virus was added to the top chamber. The amount of infectious virus in the top chamber was quantitated by plaque assay at 48 hours post infection. Three experimental replicates were performed. Data points show mean values of each experimental replicate.

### Adipocyte STING activation correlates with antiviral activity

The cGAS-STING pathway is a critical driver of the innate immune response to infection that leads to the production of antiviral cytokines^19^. This pathway is activated by the presence of cytosolic DNA, which can be produced during infection or through the degradation of cytosolic mitochondria^20, 21^. Previous studies have shown that in mice, adipocytes exhibit elevated levels of STING signaling even in the absence of infection, likely due to constitutive autophagic degradation of mitochondria^22–24^. STING activation leads to IFN production through the phosphorylation and translocation of IRF3 and IRF7 to the nucleus^25^. To determine whether our hPSC-derived adipocytes showed elevated levels of nuclear IRF3 and IRF7 relative to other hPSC-derived cell types, we quantified nuclear staining of each of these proteins. Our results showed that, relative to hPSC-derived SMCs or PCs, hPSC-derived adipocytes had significantly higher levels of nuclear IRF3 and IRF7 **(Fig. 3A-B & Fig. S6),** suggesting baseline activation of the STING signaling pathway.

**Figure 3.**
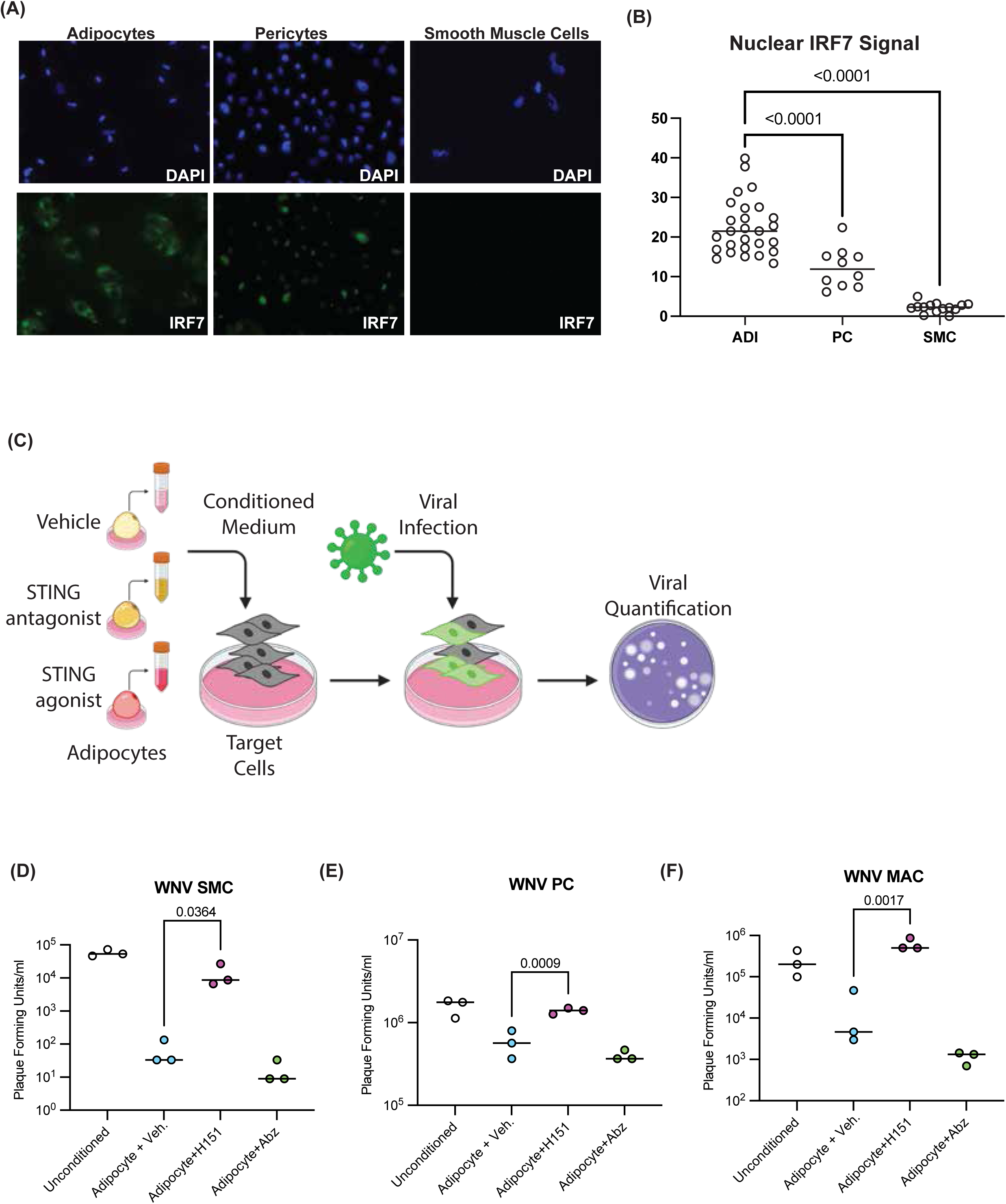
STING signaling in adipocytes is essential for generation of antiviral state in neighboring cells: **(A)** Representative images of immunofluorescence staining of IRF7 in adipocytes, pericytes, and smooth muscle cells. **(B)** Quantification of nuclear staining shown in A. A minimum of five cells were quantitated for each experimental replicate. Two experimental replicates were performed. **(C)** Schematic description of the experimental setup. **(D-F)** Plaque assays showing released virus from smooth muscle cells (D), pericytes (E), or macrophages (F) treated with unconditioned medium, or adipocyte medium +/- the STING antagonist H151 or the STING agonist diABZI prior to infection with WNV. Released infectious virus was quantitated at 48 hours post-infection. Data points show individual experimental replicates with means.

We next investigated if reducing activation of STING in adipocytes would disrupt the effect of conditioned medium on target cells. We exposed adipocytes to both STING pathway agonists and antagonists before collecting conditioned media **(Fig. 3C)**. Inhibiting STING signaling with the antagonist H151 significantly reduced the effect of adipocyte-conditioned medium on target cell viral production **(Fig. 3D-F)**. Conversely, treatment of adipocytes with the STING agonist, diABZI, resulted in a slight increase of the antiviral activity of adipocyte-conditioned medium **(Fig. 3D-F).** Notably, directly inhibiting STING activity in target cells during infection had no impact on viral replication **(Fig. S7)**. This finding is consistent with reports that flaviviruses inhibit STING activation during infection^26^. Conversely, exogenous activation of STING during infection reduced viral replication **(Fig. S7)**. Collectively, these data demonstrate that STING activity in adipocytes is essential for producing antiviral factors that, when released, protect neighboring cells from infection.

### Adipocytes activate target cell interferon signaling to confer viral protection

Activation of the STING pathway results in the production of type I interferons, including IFNα. To determine if our hPSC-derived adipocytes released more IFN-alpha relative to adipocyte progenitor cells, we measured the amount of IFNα protein in either unconditioned media, progenitor conditioned media, or adipocyte conditioned media. Our results showed that adipocytes secrete elevated levels of IFNα relative to adipocyte progenitor cells **(Fig. 4A)**. In addition, we performed RNA sequencing on SMCs treated with adipocyte- or progenitor-conditioned medium and, through gene set enrichment analysis, identified the response to interferons α and γ as the two most upregulated gene sets **(Fig. 4B)**. This robust response to treatment with adipocyte-conditioned medium was also seen via detection of differences in the expression levels for individual genes **(Fig. S8A-B)**. In contrast, progenitor-conditioned medium induced no such significant changes in gene expression **(Fig. S8C)**. The addition of IFNα to medium conditioned by adipocyte precursor cells resulted in this medium having similar antiviral properties as were observed with adipocyte conditioned medium **(Fig. 4C-D).** Lastly, inhibiting the activity of IFNα in the adipocyte-conditioned medium through the addition of a neutralizing IFNα antibody reduced the antiviral effect of adipocyte-conditioned medium on target cells (**Fig. 4E-F)**. Based on these results, we hypothesized that IFNα signaling is the pathway through which adipocytes confer viral protection. Taken together, these findings demonstrate that healthy adipocytes release factors that have a protective effect against viral infection. Additionally, the data suggest STING and IFNα signaling as the most likely axis through which the viral protection acts.

**Figure 4:**
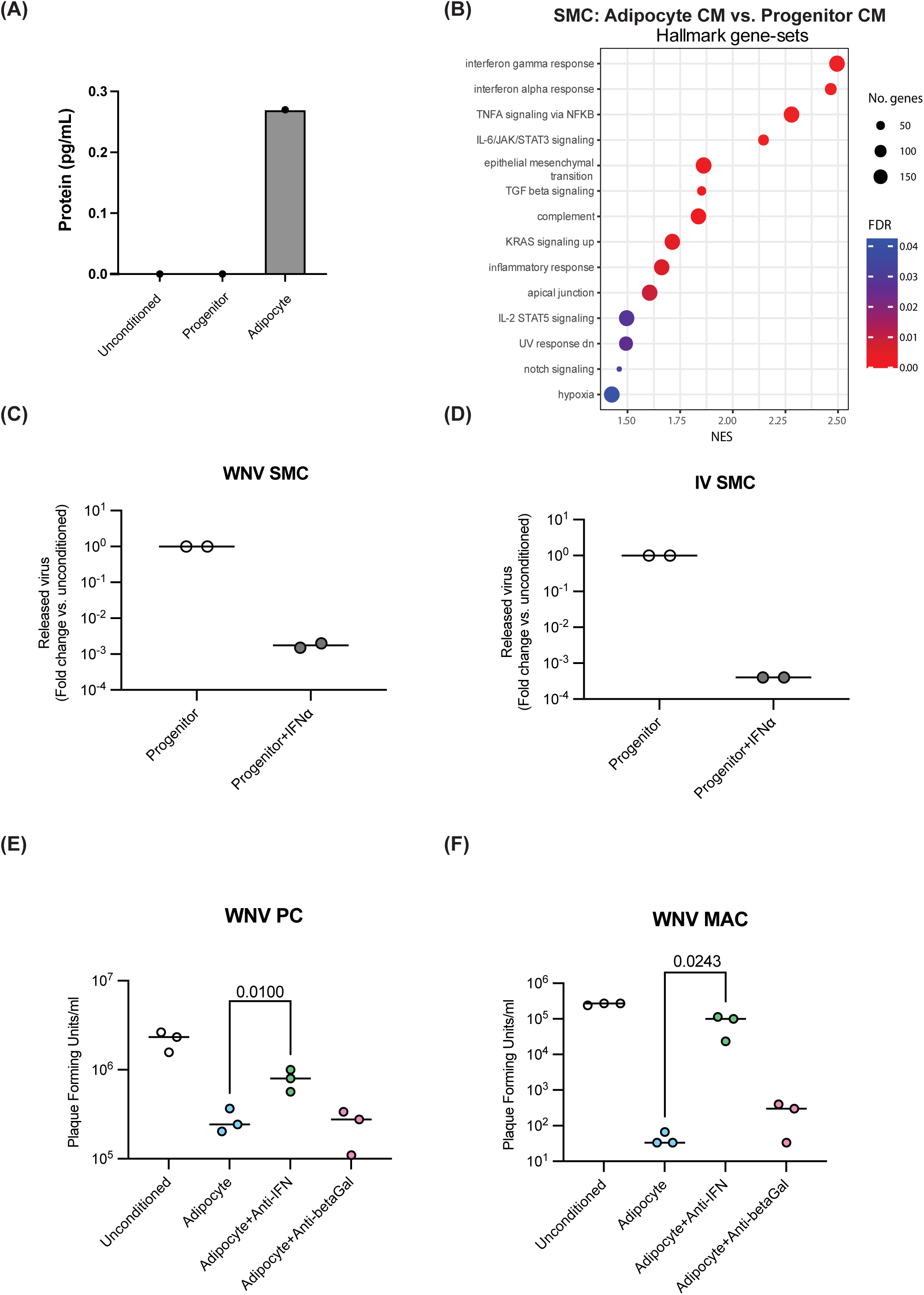
IFN signaling is critical to adipocyte antiviral activity: **(A)** Quantitation of IFNα from unconditioned adipocyte medium, progenitor-conditioned medium, or adipocyte-conditioned medium. Results are shown as mean from two experimental replicates. **(B)** Bulk RNA-sequencing was performed on total cellular RNA from smooth muscle cells (SMCs) treated with adipocyte-conditioned medium or progenitor conditioned medium. Dot plot shows gene-sets from the Hallmark collection^38^of the MSigDB that were enriched (FDR < 0.05) using gene-set enrichment analysis (GSEA)^39^. **(C,D)** Plaque assay quantification of virus released from WNV infected SMCs. SMCs were treated with adipocyte progenitor conditioned medium with or without the addition of IFNα for 24 hours prior to infection with WNV. **(E,F)** Plaque assay quantification of released virus from pericytes (E), or macrophages (F) treated with unconditioned medium, or adipocyte-conditioned medium +/- an anti-IFNα antibody or control (Beta-Gal) antibody prior to infection with WNV. Released infectious virus was quantitated at 48 hours post-infection. Data points show individual experimental replicates with means.

### Treatments that disrupt adipocyte homeostasis reduce adipocyte antiviral properties

Viral infections often have worse outcomes in individuals with obesity or lipodystrophy; conditions associated with dysregulated adipocytes^27–30^. To investigate whether the antiviral activity of adipocytes is influenced by metabolic health, we examined the effects of dexamethasone, an agent known to disrupt lipid homeostasis and adipocyte metabolism^31^, on the antiviral activity of adipocytes. We exposed adipocytes to dexamethasone for five days, at which point the dexamethasone-containing medium was removed and replaced with fresh medium without dexamethasone. Adipocyte-conditioned medium was harvested 48 hours after dexamethasone was removed **(Fig.5A)**. Similar to our observations using medium from adipocytes treated with a STING antagonist, medium from dexamethasone-treated adipocytes showed a reduced antiviral effect on SMCs, PCs, and MACs **(Fig. 5B-D)**.

**Figure 5.**
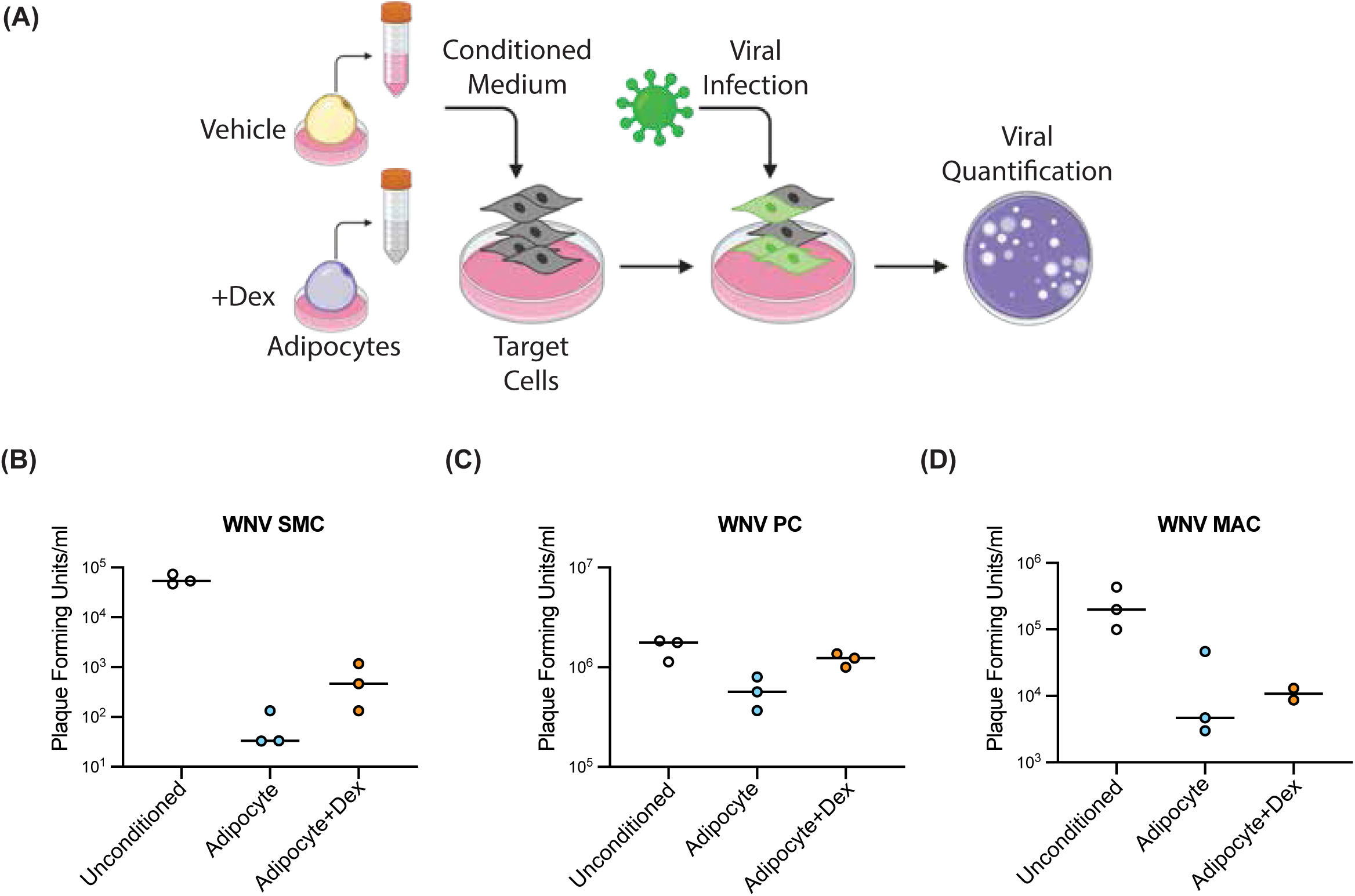
Pharmacological modulation of adipocyte antiviral activity: **(A**) Schematic description of the experimental setup. Conditioned medium was collected from adipocytes with or without dexamethasone (Dex) pretreatment. **(B-D)** Plaque assay quantitation of released virus from smooth muscle cells (B), pericytes (C), or macrophages (D) treated with unconditioned, adipocyte-conditioned, or adipocyte+Dex- conditioned medium prior to West Nile Virus (WNV) infection. Released virus was quantitated at 48 hours post-infection. Data points show individual experimental replicates with means.

Finally, we examined the release of cytokines with known immunomodulatory properties in addition to IFNα. We employed a cytokine array to determine the levels of various immune-signaling proteins released from adipocytes. A number of cytokines were elevated in adipocyte-conditioned medium compared to unconditioned or progenitor- conditioned medium **(Fig. S9A)**. We prioritized factors that were also downregulated by dexamethasone treatment, given our results showing that dexamethasone treatment reduced adipocyte antiviral activity. Critically, IFNα was one of the factors that was reduced with dexamethasone treatment. In addition, two other proteins in our cytokine panel met the criteria: IL-15 and CD40L **(Fig. S9A)**. To determine if any of these candidate cytokines were sufficient to induce the antiviral effects observed with adipocyte-conditioned medium, we added each candidate protein to progenitor- conditioned medium and then treated target cells with the modified medium. Our results show that of our three candidate proteins, only the addition of IFNα recapitulated the antiviral activity observed in adipocyte-conditioned medium **(Fig S9B-C)**.

## Discussion

This study demonstrates a novel role for adipocytes in reducing the infectivity of nearby cells through STING-dependent release of an antiviral factor or factors. Our investigation suggests that one of these factors is IFNα. Our findings challenge the traditional view of adipocytes as solely energy storage cells, highlighting their active contribution to the innate immune response against viral infections.

It has been shown that metabolically unhealthy conditions, including obesity, are associated with chronic low-grade inflammation of adipose tissue and increased susceptibility to viral infections, likely due to dysregulated adipocytes influencing the local immune milieu^32–34^. Our study provides a potential mechanism for this observation by demonstrating that metabolically healthy adipocytes can reduce the production of infectious virus in several cell types through the secretion of IFNα. This finding highlights the significance of metabolic health in supporting an effective antiviral immune response.

Our study focused on flaviviruses, which are emerging as a global health threat due to their increasing prevalence and potential to cause severe diseases. We found that adipocyte-secreted factors can protect multiple cell types from infection by a panel of flaviviruses, including West Nile virus, Zika virus, and Ilheus virus. In addition, adipocyte-secreted factors also protect smooth muscle cells from SARS-CoV-2 infection. This broad protective effect suggests that adipocytes may play a crucial role in the host defense against a range of viral pathogens. Furthermore, we showed that pharmacological inhibition of the STING pathway in adipocytes abrogated their antiviral activity. Consistent with previous findings in mouse adipocytes^35^, we show that healthy hPSC-derived adipocytes exhibit constitutive activation of the STING pathway, which likely contributes to higher baseline levels of IFNα secretion. This constitutive activation may, in part, explain why adipocytes have lower levels of infectious virus production relative to vascular and immune cells **(Fig. S1 & Fig. 2)**. We hypothesize that this constitutive low level of inflammatory signaling in adipocytes may serve as an endogenous mechanism to reduce the spread of infection in the vasculature and circulating immune cells. We also observed that expression of the gene encoding for TXNIP, thioredoxin-interacting protein, is significantly down-regulated in SMCs following treatment with adipocyte-conditioned media **(Fig. S8**). A recent study showed that knockdown of TXNIP is antiviral against HSV-1^36^. It is possible that the reduction in TXNIP expression following exposure to adipocyte-secreted factors contributes to the antiviral state observed in our target cells.

Our results show that the antiviral effect of adipocyte-conditioned medium does not extend to endothelial cells, which was surprising given that endothelial cells are known to respond to IFNα. To the best of our knowledge, there have been no direct comparisons of the sensitivity of different vascular cells to IFNα, raising the possibility that endothelial cells are less sensitive than mural cells to IFNα exposure. Notably, since IFN-alpha is known to cause endothelial cell dysfunction, a reduced sensitivity to IFNα may be protective. Future experiments will assess the sensitivity of different vascular cell populations to low levels of IFNα exposure. Additionally, adipocytes may secrete other antiviral factors to which endothelial cells do not respond.

While our study provides compelling evidence for the antiviral role of adipocytes, it is limited in its use of only *in vitro* model systems, which may not fully recapitulate the complexity of in vivo interactions. However, our results are in part supported by data in a mouse model of alphavirus infection, which showed that although obese mice had increased morbidity following infection, the amount of infectious virus in the blood was significantly reduced relative to lean mice^37^. Our future studies will investigate the role of adipocytes in antiviral immunity in vivo using animal models of viral infection. Additionally, our study focused primarily on flaviviruses, and further research is needed to determine whether adipocytes play a similar role in the defense against other viral pathogens.

Collectively, our study provides novel insights into the interplay between metabolism and immunity, highlighting the importance of adipocytes in shaping the antiviral immune response. These findings have implications for understanding the pathogenesis of viral infections and may inform the development of novel therapeutic strategies targeting adipocyte function to enhance antiviral immunity.

## Acknowledgements

We thank the Genome Technology Core and Keck Imaging Facility at Whitehead Institute for Biomedical Research. This work was supported by NIH grant U19AI131135-01, NIBIB grant T32 EB016652, and by a generous gift from Jim Stone. The following reagent was deposited by the Centers for Disease Control and Prevention and obtained through BEI Resources, NIAID, NIH: SARS-Related Coronavirus 2, Isolate USA-WA1/2020, NR-52281. This work was performed in part in the Ragon Institute BSL3 core, which is supported by the NIH-funded Harvard University Center for AIDS Research (P30 AI060354). Some of the figure panels were made with biorender.com.

---

**Figure S1.**
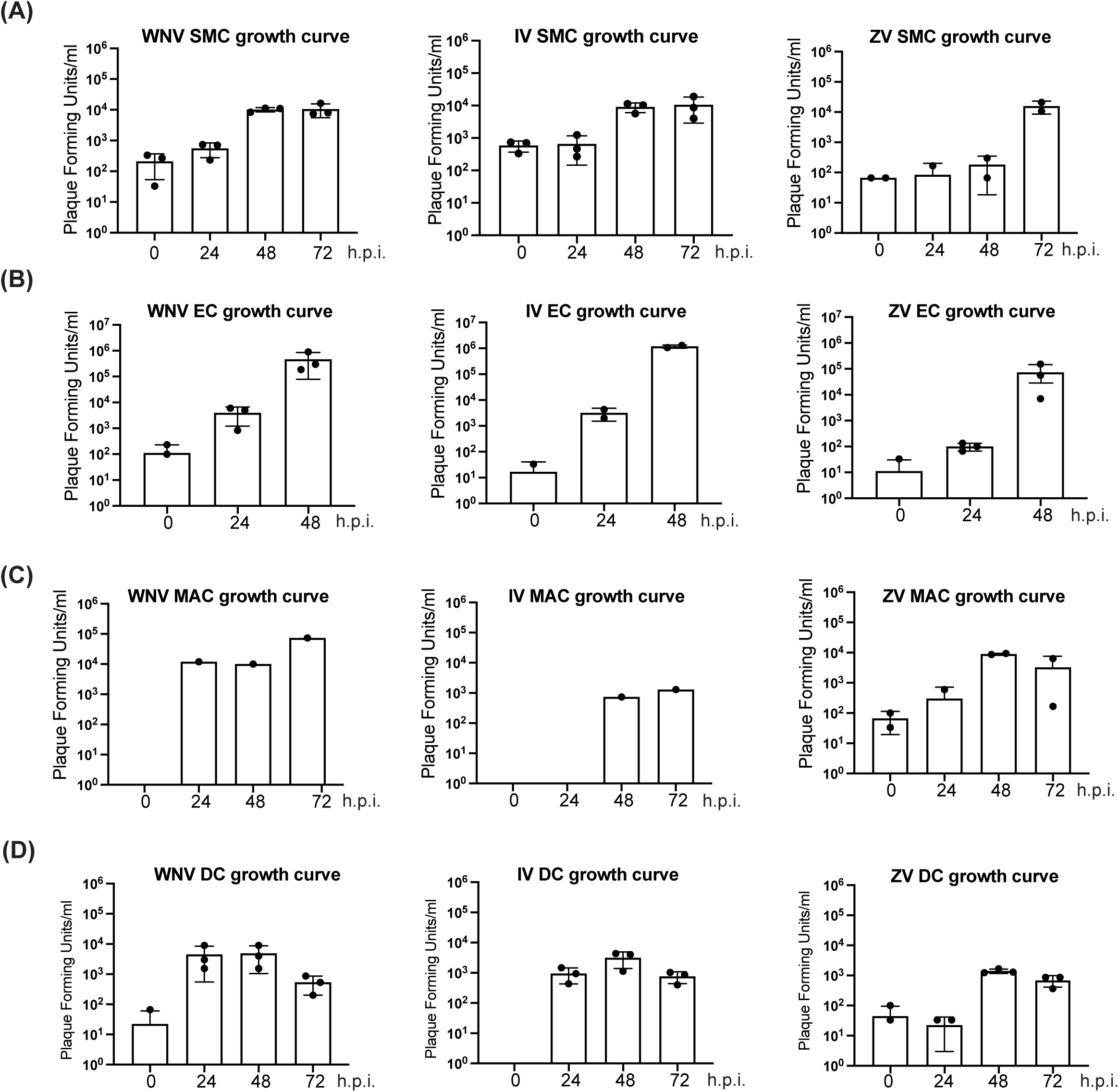
Replication kinetics of flaviviruses in vascular cells, immune cells, and adipocytes: **(A-D)** Plaque assay quantitation of released infectious virus from hPSC-derived smooth muscle cells (A), endothelial cells (B), macrophages (C), dendritic cells (D) at the indicated hour post infection (h.p.i.) with West Nile Virus (left column), Ilheus Virus (middle column), or Zika Virus (right column). Data points show individual replicates with mean bar graphs, and standard deviation error bars.

**Figure S2.**
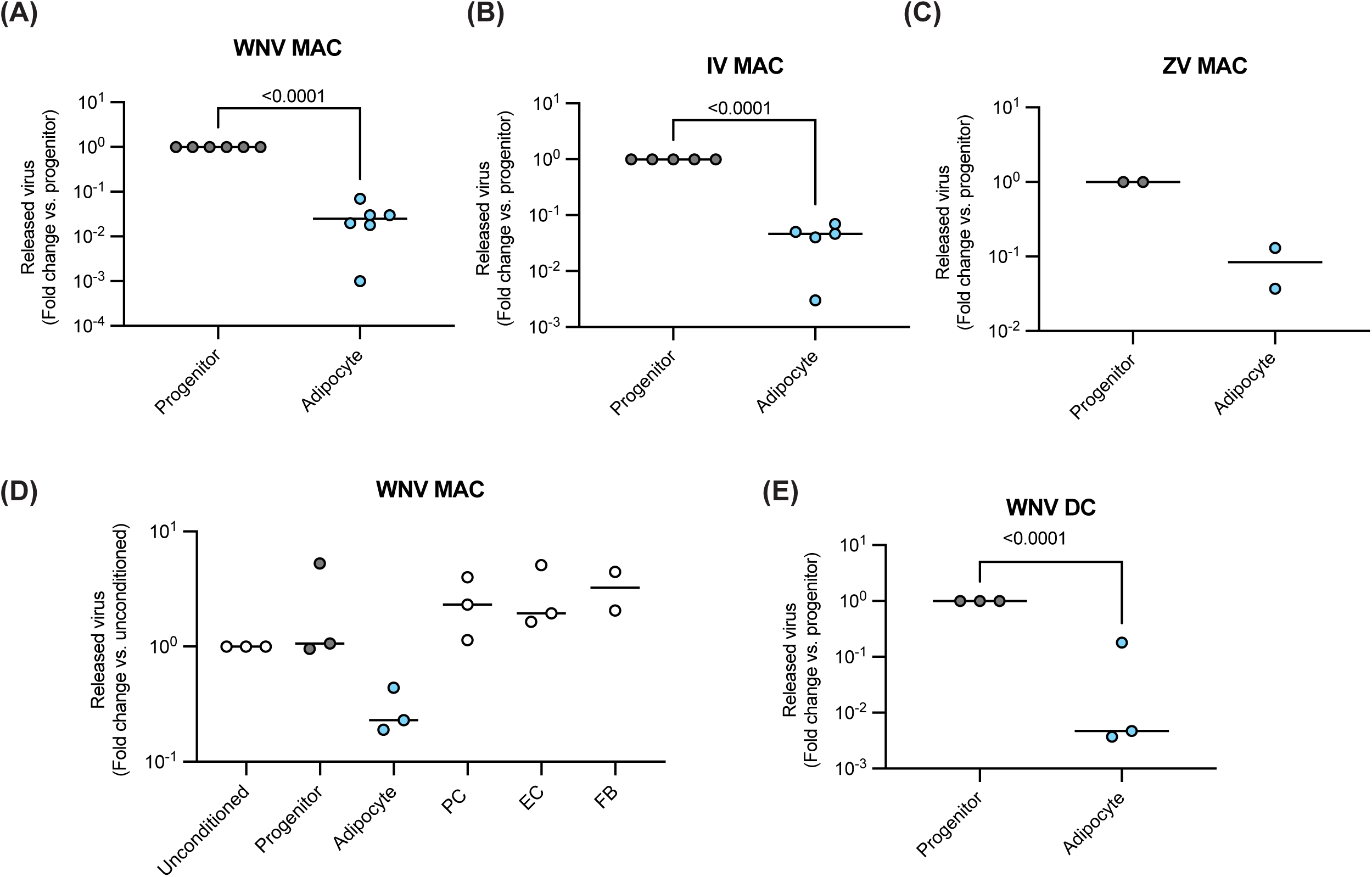
Adipocytes secrete factors that restrict infection of immune cells. **(A-C)** Plaque assays showing released virus from hPSC-derived macrophages treated with progenitor-conditioned or adipocyte-conditioned medium prior to infection with West Nile Virus (A), Ilheus Virus (B), or Zika Virus (C). **(D)** Plaque assay quantitation of released virus from macrophages treated with unconditioned, progenitor-, adipocyte-, pericyte-, endothelial cell-, or fibroblast-conditioned medium prior to infection with West Nile virus. **(E)** Plaque assays showing released virus from dendritic cells treated with progenitor-conditioned or adipocyte-conditioned medium prior to infection with West Nile Virus. Data points show individual experimental replicates with means.

**Figure S3.**
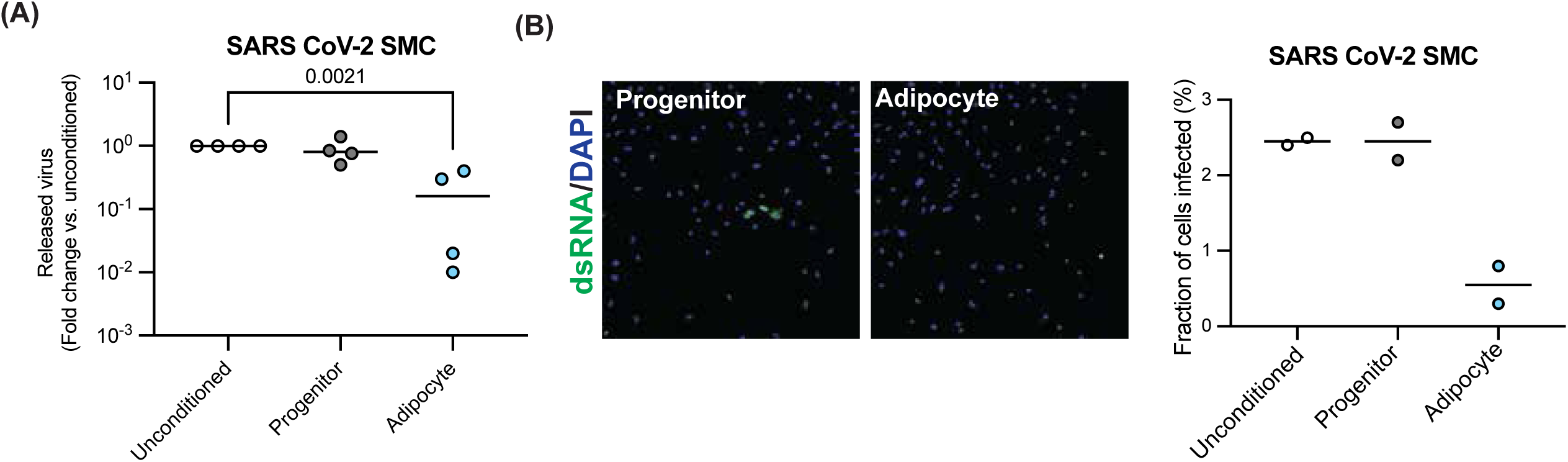
**The protective effect of adipocyte conditioned medium extends beyond flaviviruses:** (A) Plaque assay quantitation of released infectious virus from smooth muscle cells treated with unconditioned, progenitor-conditioned or adipocyte-conditioned medium prior to infection with SARS-CoV-2. Released virus was quantitated at 48 hours post-infection. **(B)** Representative images of immunofluorescence staining of dsRNA in smooth muscle cells treated with unconditioned, progenitor-conditioned or adipocyte- conditioned medium prior infection with SARS-CoV-2. Cells were fixed and stained at 48 hours post-infection. **(C)** Quantification of staining shown in B. Data points show individual replicates with means.

**Figure S4.**
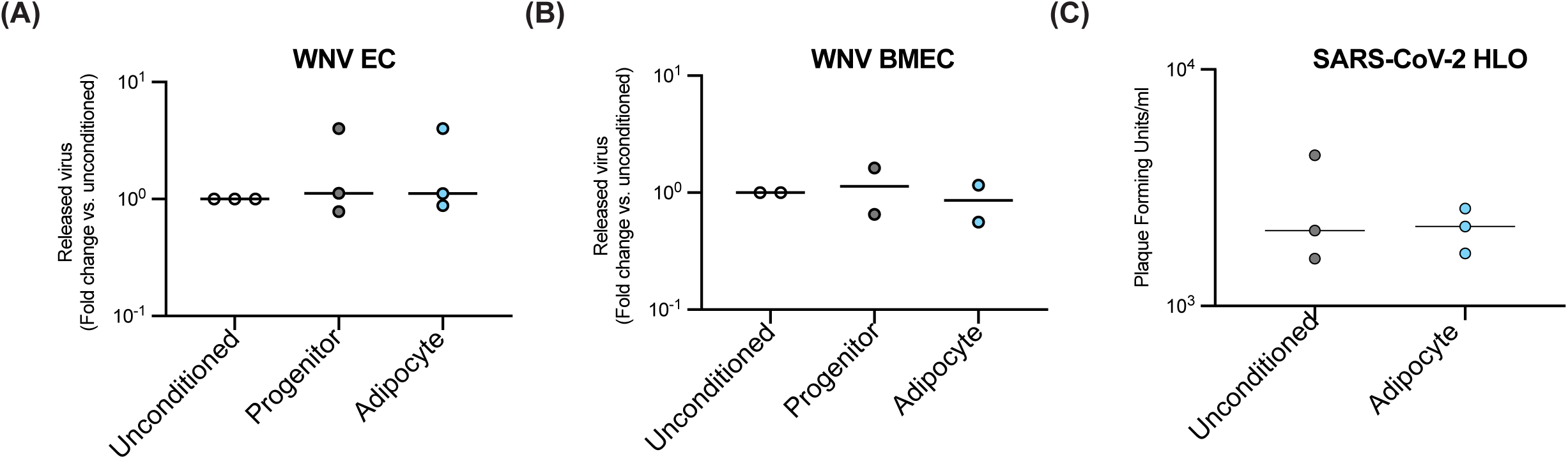
Adipocyte conditioned medium does not protect all cell types: **(A,B)** Plaque assay quantitation of released infectious virus from hPSC-derived endothelial cells (EC) (A) or hPSC-derived brain microvascular endothelial cells (BMEC) (B) treated with unconditioned, progenitor-conditioned or adipocyte-conditioned medium prior to infection with West Nile virus. Released infectious virus was quantitated at 48 hours post-infection. **(C)** Plaque assay quantitation of released infectious virus from hPSC-derived liver organoids treated with unconditioned, progenitor-conditioned or adipocyte-conditioned medium prior to infection with SARS-CoV-2. Released virus was quantitated at 48 hours post-infection. Data points show individual experimental replicates with means.

**Figure S5.**
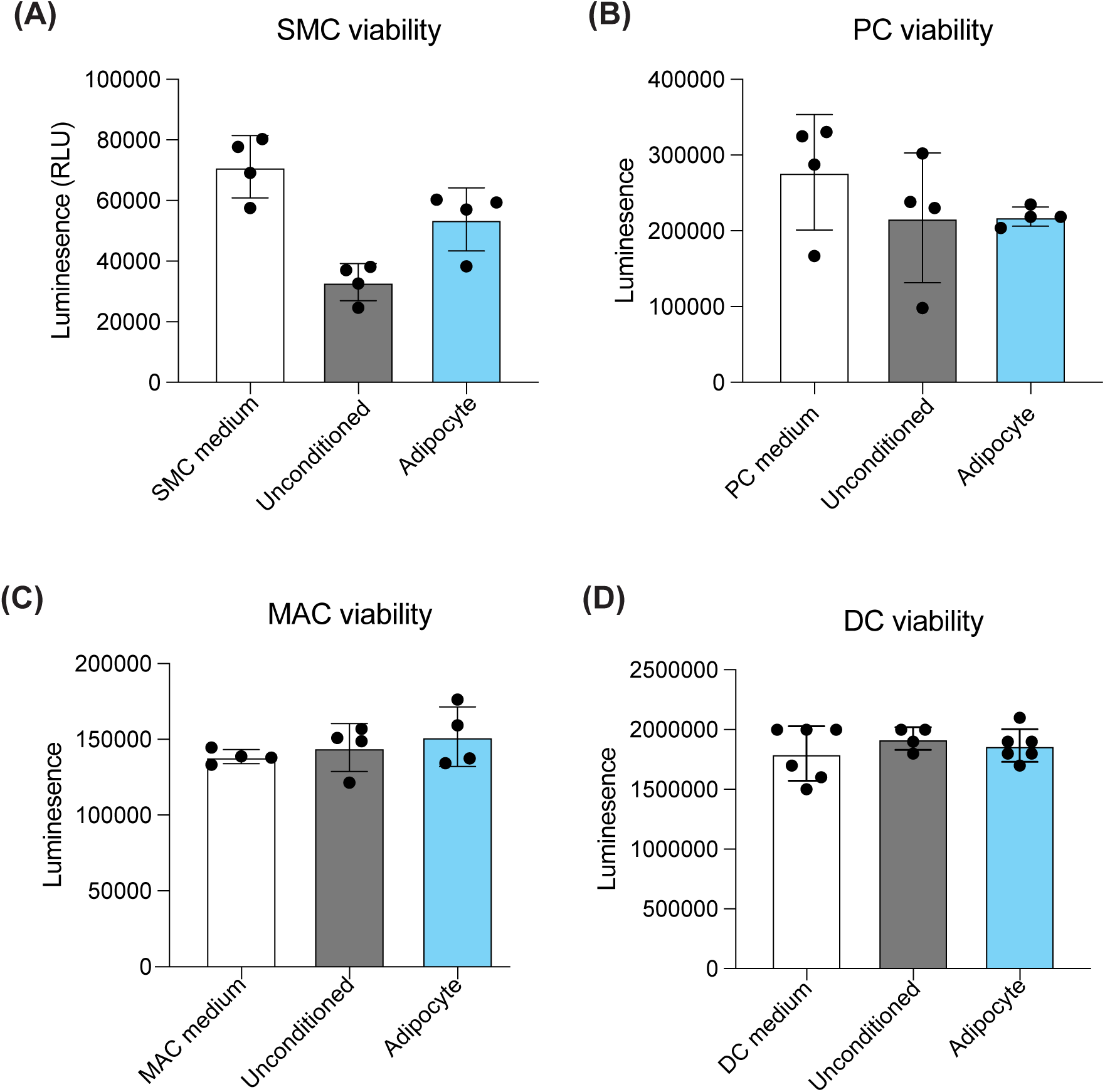
Adipocyte conditioned medium does not reduce cell viability: **(A-D)** Cell Titer Glo viability assays were performed on hPSC-derived smooth muscle cells (SMC) (A), hPSC-derived pericytes (PC) (B), hPSC-derived macrophages (MAC) (C), or hPSC-derived dendritic cells (DC) (D) after 72 hours of culture in either cell-type specific culture medium, unconditioned adipocyte medium, or adipocyte-conditioned medium. Data points show individual replicates with mean bar graphs, and standard deviation error bars.

**Figure S6.**
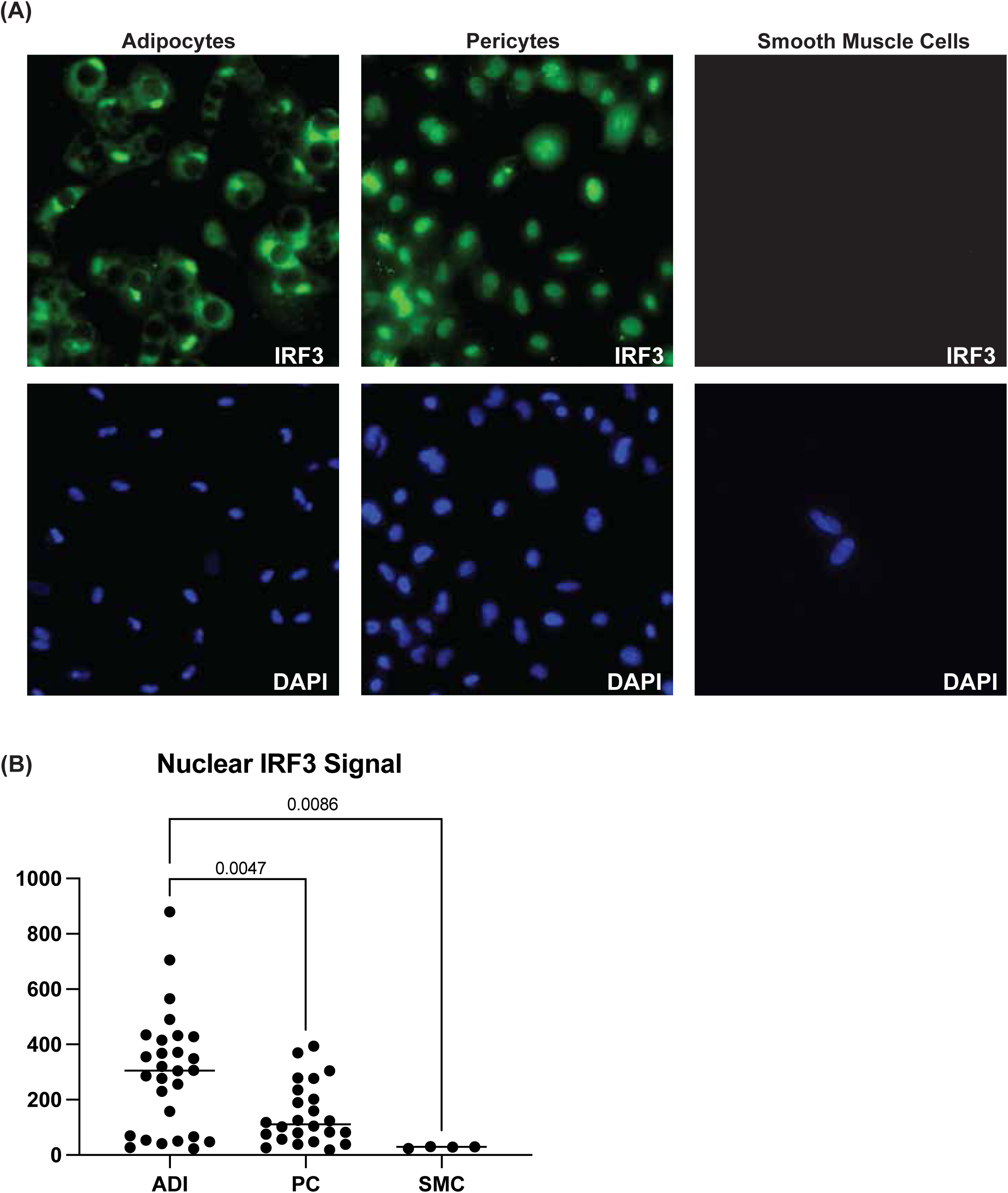
Adipocytes display higher IRF3 activation: **(A)** Representative images of immunofluorescence staining of IRF3 in adipocytes, smooth muscle cells, and pericytes. **(B)** Quantification of nuclear staining shown in (A). For adipocytes and pericytes a minimum of 10 cells were quantitated for each experimental replicate. For smooth muscle cells two cells were quantitated for each experimental replicate. Two experimental replicates were performed for all cell types.

**Figure S7.**
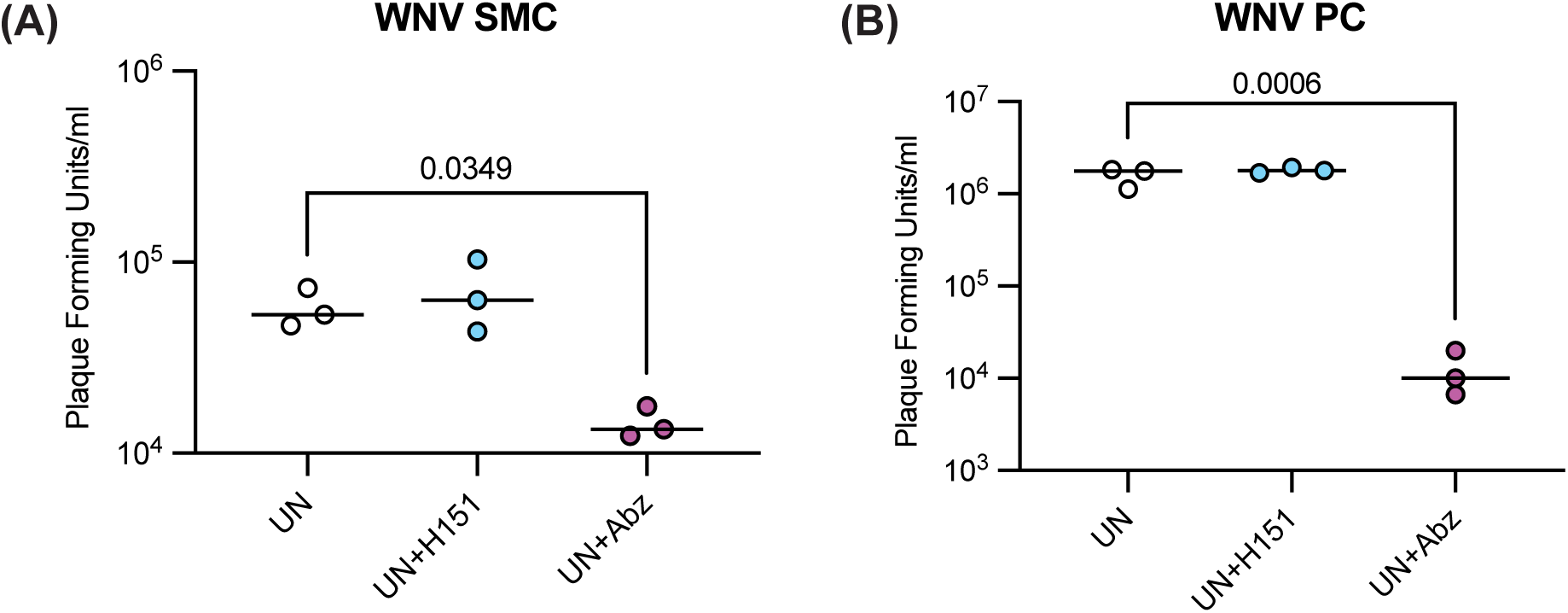
Inhibition of STING activity in infected target cells does not reduce viral infection: **(A,B)** Plaque assay quantitation of infectious virus released from smooth muscle cells (A) or pericytes (B) with or without treatment with the STING agonist diABZI or the STING antagonist H151, or dexamethasone (Dex) prior to infection with West Nile Virus. Released infectious virus was quantitated at 48 hours post-infection. Data points show individual replicates with means.

**Figure S8.**
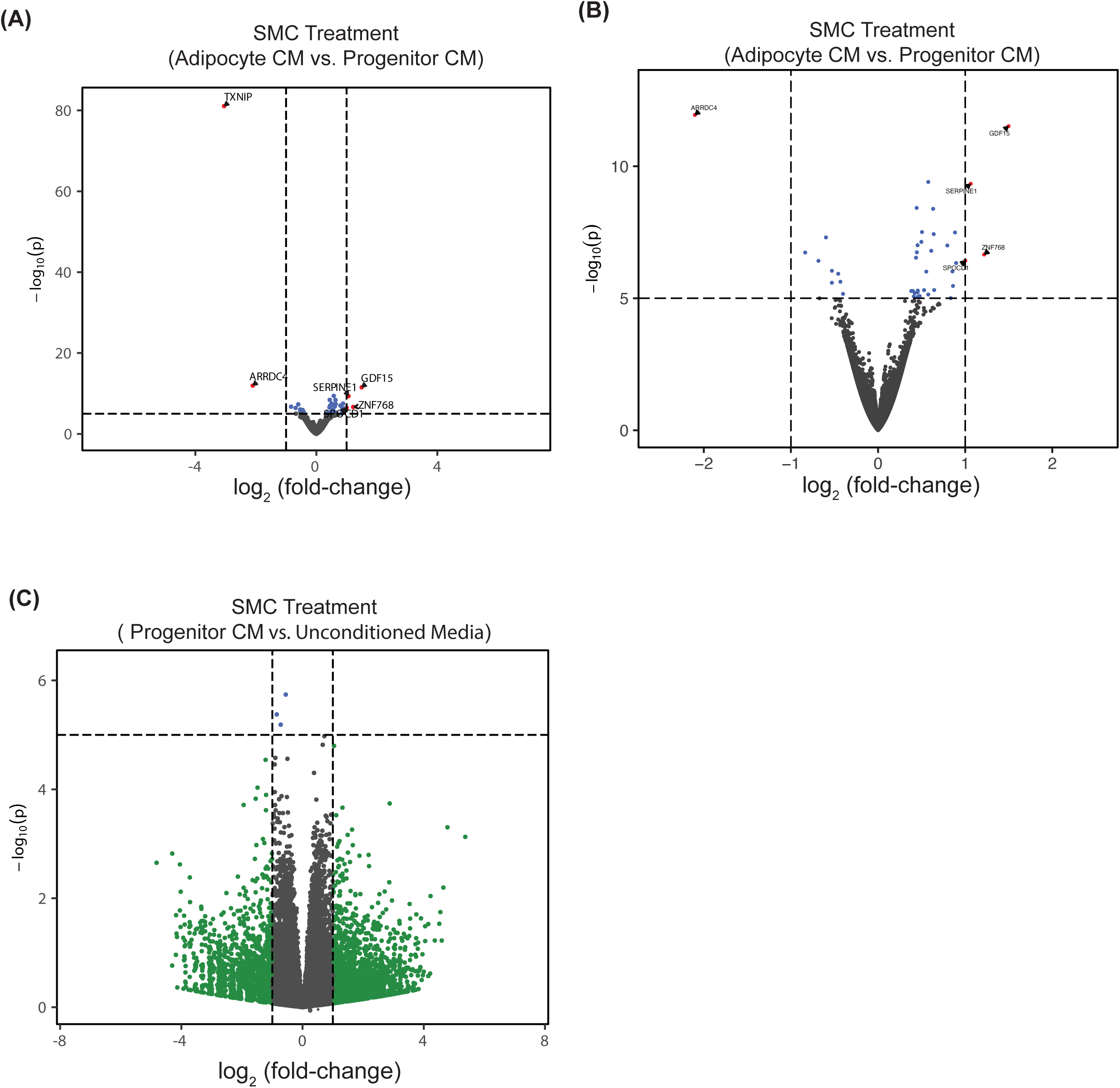
Treatment with adipocyte conditioned medium induces transcriptional changes in SMCs: **(A-C)** Bulk RNA-sequencing was performed on total cellular RNA from SMCs treated with adipocyte-conditioned medium or progenitor conditioned medium. (A) Volcano plot showing differentially expressed genes in SMCs treated adipocyte-conditioned medium compared to SMCs treated with progenitor-conditioned medium. (B) Volcano plot as in A, but zoomed in for visual clarity in the absence of the TXNIP gene. (C) Volcano plot showing differentially expressed genes in SMCs treated progenitor-conditioned medium compared to SMCs treated with unconditioned medium.

**Figure S9.**
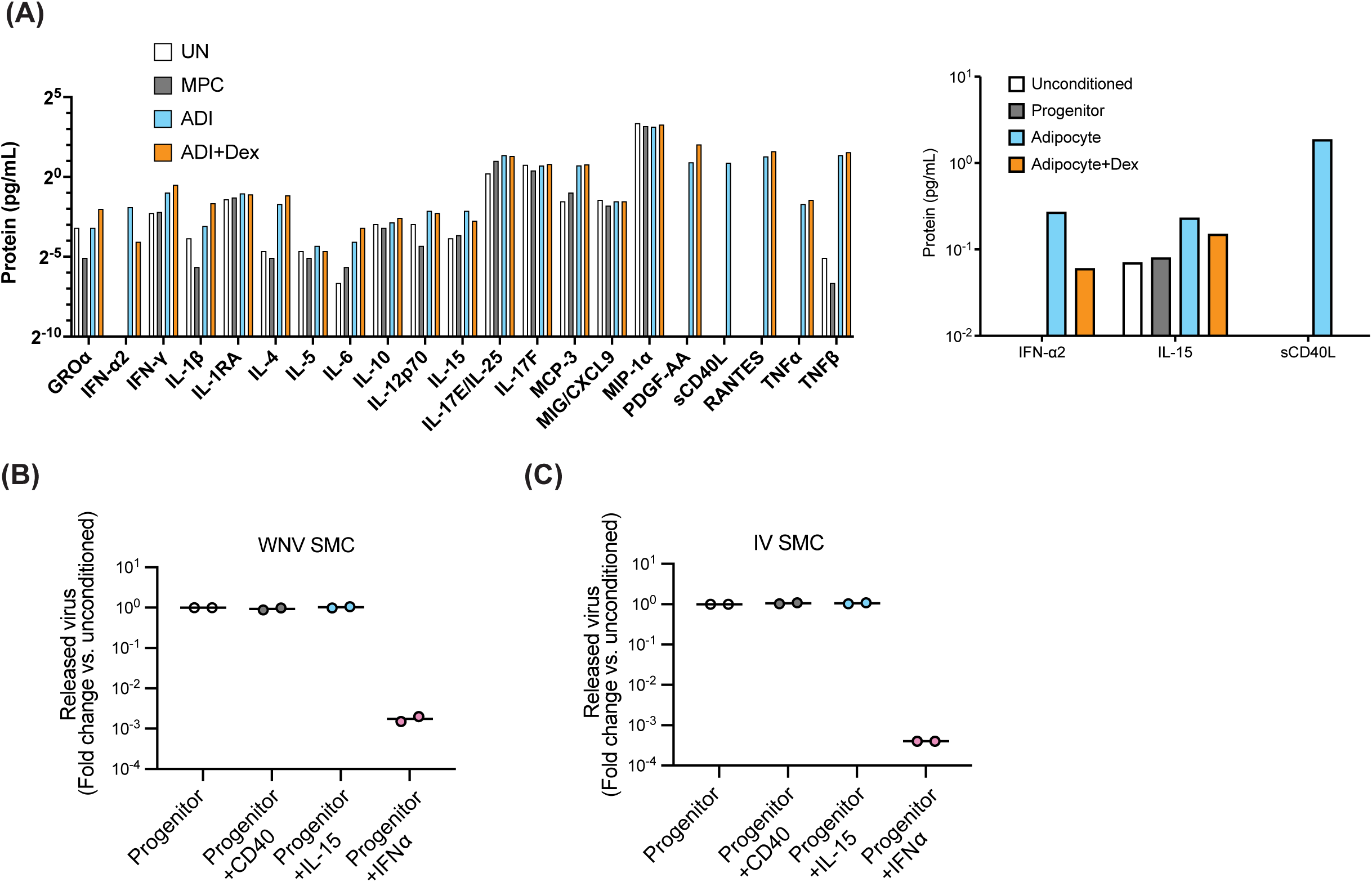
Testing additional potential antiviral adipocyte secreted factors: **(A)** Quantitation of selected cytokines in unconditioned-medium, progenitor-conditioned medium, adipocyte-conditioned medium, and adipocyte+Dex- conditioned medium. The right panel is a zoomed in subset of the three selected candidates. **(B,C)** SMCs were treated with progenitor-conditioned medium with or without the addition of the selected cytokines shown in (A) prior to infection with West Nile virus (B) or Ilheus virus (C). Released infectious virus was quantitated at 48 hours post infection. Released virus was quantitated at 48 hours-post-infection. Data points show individual experimental replicates with means.

## Methods

### hPSC culture

H1 (WA01) human pluripotent stem cells (hPSCs) were obtained from WiCell. hPSCs were maintained feeder-free on Matrigel (Corning, 354234) in StemFlex medium (Thermo Fisher Scientific, A3349401) and passaged as clumps using Versene solution (Thermo Fisher Scientific, 15040066).

### Adipocyte differentiation

Differentiation of hPSCs into adipocytes was performed as previously published^40^. Briefly, hPSCs were passaged and grown in suspension culture to form embryoid bodies. After 1 week, the embryoid bodies were plated on tissue culture–treated plastic and grown in DMEM + 10% fetal bovine serum. The outgrowing mesenchymal progenitor cells (MPCs) were used from passage 4 through 8. For adipocyte differentiation, MPCs were infected with previously described lentivirus to express the transactivator reverse tetracycline-controlled transactivator (rtTA) and inducible expression of peroxisome proliferator-activated receptor gamma isoform-2 (PPARg2). MPCs were passaged and grown to confluence and then exposed to doxycycline (700 ng/ml) for 2 weeks in previously established A2 medium and then 1 week without doxycycline in A2 medium. Afterward, adipocytes were exposed to the experimental treatment medium for 5 days unless otherwise indicated.

### Smooth muscle cell differentiation

Smooth muscle cells were differentiated as described previously^41^. Briefly, H1 hPSCs were cultured in E8 medium (Thermo Fisher Scientific) and dissociated using Accutase (Thermo Fisher Scientific) into a single cell suspension and plated at 15,000 cells/cm^2^ in E8 medium with Y-27632 (10M). Y-27632 was removed after 24 hours and cells cultured in E8 medium until they reach approximately 70% confluency at which point the medium was replaced with MelM media, this is considered day 0. On day 2 the medium was replaced with PC/SMC1 media. On day 5 the medium was replaced with PC/SMC2 medium for 24 hours and then cells were passaged 1:3 on to fibronectin (FC01010MG, Fisher Scientific) coated tissue culture plates in SMC3 media. Plates were coated with fibronectin at 10 ug/ml for 30 minutes at 37 C prior to the addition of cells. After 48 hours the medium was replaced with SMC4 media. After 48 hours the medium was replaced with SMC5 and cells kept in culture for 7 days prior to use. All medium formulations can be found in (Richards et al., 2024)^41^.

### Pericyte differentiation

Pericytes were differentiated as described previously^41^. Briefly, H1 hPSCs were cultured in E8 medium (Thermo Fisher Scientific) and dissociated using Accutase (Thermo Fisher Scientific) into a single cell suspension and plated at 15,000 cells/cm^2^ in E8 medium with Y-27632 (10M). Y-27632 was removed after 24 hours and cells cultured in E8 medium until they reach approximately 70% confluency at which point the medium was replaced with MelM media, this is considered day 0. On day 2 the medium was replaced with PC/SMC1 media. On day 5 the medium was replaced with PC/SMC2 medium for 24 hours and then cells were passaged 1:3 on to fibronectin (FC01010MG, Fisher Scientific) coated tissue culture plates in PC3 media. Plates were coated with fibronectin at 10 ug/ml for 30 minutes at 37 C prior to the addition of cells. After 48 hours the medium was replaced with PC4 media. After 48 hours the medium was replaced with fresh PC4 and cells kept in culture for 7 days prior to use. All medium formulations can be found in (Richards et al., 2024)^41^.

### Macrophage differentiation

HMacs were derived as previously described^18^. Briefly, myeloid progenitor cells were derived from H1 embryonic stem cells (ESC) as previously described (Brownjohn et al., 2018). Briefly, H1 ESCs were cultured and maintained in feeder-free culture conditions in StemFlex (Gibco) media. At approximately 70% confluence, the medium was transitioned to Essential 8 (ThermoFisher Scientific) pluripotent stem cell medium supplemented with P/S and 10 µM Y-27632 (ROCK) inhibitor (Stem Cell Technologies) on day -1. On day 0, the H1 ESCs were dissociated into a single cell suspension using TrypLE Express (ThermoFisher Scientific) and diluted to 0.666x10^6^ per mL in embryoid body (EB) medium consisting of complete E8 medium with P/S supplemented 50 ng/mL bone morphogenetic protein 4 (Peprotech), 20 mg/mL stem cell factor (Peprotech), and 50 ng/mL vascular endothelial growth factor, with 10 µM ROCK inhibitor. 150 µL of this cell suspension per well was plated into a 96-well U-bottom ultra-low adherence well plates (Corning) and centrifuged at 200 RCF for 5 min. On day 2, 150 µL of EB medium without ROCK inhibitor was added to each well. On day 4, the EBs were transferred to a 10 cm tissue culture-treated polystyrene dish in 12 mL of hematopoetic myeloid (HpM) medium consisting of X-VIVO 15 medium (Lonza) supplemented with 2 mM GlutaMax (Gibco), 55 µM beta-mercaptoethanol (Sigma Aldrich), 100 ng/mL macrophage colony-stimulating factor (Peprotech), and 25 ng/mL interleukin-3 (Peprotech). Medium was replaced with fresh HM medium every 3-4 days. At 2-3 weeks after EB plating, floating CD14+ myeloid precursors (MPs) were collected and differentiated into macrophages by plating the cells in a 10 cm tissue culture-treated polystyrene dish in 12 mL serum-free macrophage medium (SFMM) consisting of glutamine containing no glucose RPMI medium (Gibco) supplemented with 1 mM glucose (Sigma Aldrich), 0.432 mg/L zinc sulfate heptahydrate (Sigma Aldrich), 7.5 mg/L human transferrin (holo) (Sigma Aldrich), 1.9 mg/L ethanolamine (Sigma Aldrich), 0.005 mg/L sodium selenite (Sigma Aldrich), 1.7 nM human insulin (Sigma Aldrich), 1:100 dilution of minimal essential amino acids (Gibco), 1:100 dilution of sodium pyruvate (Gibco), P/S, 0.2 g/L of recombinant human albumin (CellaStim), and 100 ng/mL macrophage colony-stimulating factor. The MPs were differentiated into macrophages over 10 days in SFFM with the same feeding schedule as the EBs in HpM medium.

### Dendritic cell differentiation

Myeloid progenitor cells were derived from H1 embryonic stem cells (ESC) as described in the previous section (Macrophage differentiation). At 2-3 weeks after EB plating, floating CD14+ myeloid precursors (MPs) were collected and differentiated into dendritic cells by plating in a 10 cm tissue culture-treated polystyrene dish in 12 mL dendritic medium (DM) consisting of aMEM-nucleosides,+glutamax (Gibco) supplemented with 10% fetal bovine serum(Gibco), 1:100 dilution of minimal essential amino acids (Gibco), 1:100 dilution of sodium pyruvate (Gibco), 1:100 dilution of Pen/Strep, 55 uM 2- mercaptoethanol,100 ng/mL hGM-CSF, 100 ng/mL hFlt-3 ligand, and 50 ng/mL hIL4.

The MPs were differentiated into dendritic cells over 10 days in DM with the same feeding schedule as the EBs in HpM medium.

### Endothelial cell differentiation

Endothelial cells were differentiated as described previously^41^. Briefly, H1 hPSCs were cultured in E8 medium (Thermo Fisher Scientific) and dissociated using Accutase (Thermo Fisher Scientific) into a single cell suspension and plated at 15,000 cells/cm^2^ in E8 medium with Y-27632 (10M) and onto Matrigel-coated tissue culture plates. Y-27632 was removed after 24 hours, and cells were cultured in E8 medium until they reached approximately 70% confluency, at which point the medium was replaced with MelM media; this is considered day 0. On day 2, the medium was replaced with EC1 media. On day 5, the medium was replaced with EC2 medium for 24 hours, and then cells were passaged 1:1 onto fibronectin (FC01010MG, Fisher Scientific) coated tissue culture plates in EC3 media. Plates were coated with fibronectin at 10 ug/ml for 30 minutes at 37 C prior to the addition of cells. After 48 hours, medium was replaced with EC4 medium for 5 days and then sorted for VE-Cadherin and PECAM1 double-positive cells using FACS. Sorted cells were plated back onto fibronectin and cultured for an additional 5 days in EC4 before cryopreservation or extended expansion in EC5 medium. All medium formulations can be found in (Richards et al., 2024)^41^.

### Brain microvascular endothelial cell differentiation

Brain microvascular endothelial cells were differentiated as described previously^42^. Briefly, H1 hPSCs were cultured in E8 medium (Thermo Fisher Scientific) and dissociated using Accutase (Thermo Fisher Scientific) into a single cell suspension and plated at 15,000 cells/cm^2^ in StemFlex medium with Y-27632 (10M). Y-27632 was removed after 24 hours and at 48 hours post-plating the medium was replaced with Unconditioned Medium (UM) (100ml Knock-out Serum Replacement (Thermo Fisher Scientific), 5ml non-essential ammino acids (Thermo Fisher Scientific), 2.5ml GlutaMax (Thermo Fisher Scientific), 5ml Pen/Strep (Thermo Fisher Scientific), 3.5l -mercapto- ethanol (Sigma), and 392.5ml DMEM/F12(1:1)), this was considered day zero. The medium was replaced daily with fresh UM. On day six the medium was changed to hESFM (Thermo Fisher Scientific) supplemented with 20 ng/mL bFGF (Peprotech), 10 µM retinoic acid (RA) (Sigma), and 1:50 B27 (Thermo Fisher Scientific). The medium was not changed for 48 hours. On day eight, cells were washed once with DPBS and incubated with Accutase for 30 minutes at 37C. Cells were collected via centrifugation and plated onto either standard tissue culture plates or transwell plates (Corning #3460). Plates were coated with 400 µg/mL collagen IV (Sigma Aldrich) and 100 µg/mL fibronectin (Fisher Scientific) overnight with collagen and fibronectin and washed 1X with PBS prior to the addition of cells. Cells were plated at a density of 250,000 cells/cm^2^, whereas for transwell plates, cells were plated at a density of 1.1 x 10^6^ cells/cm2 on the transwell membrane. bFGF and RA were removed from the medium 24 hours after plating.

### Liver organoid differentiation

The HLO differentiation protocol is adapted from a previously published protocol^18^. Briefly, hPSCs were detached and seeded on Matrigel coated tissue culture plates. Medium was changed to RPMI 1640 medium containing 100 ng/mL Activin A and 50 ng/mL bone morphogenetic protein 4 (BMP4) on day 1, 100 ng/mL Activin A and 0.2% fetal bovine serum (FBS) on day 2, and 100 ng/mL Activin A and 2% FBS on day 3. On day 4–8, cells were cultured in Advanced DMEM/F12 with 500 ng/mL fibroblast growth factor 4 and 3uM CHIR99021. On day 9, cells were gently pipetted to delaminate from the dish, centrifuged at 800rpm for 3 minutes and resuspended at 1 × 106 cells/mL in Advanced DMEM/F12 with 2uM retinoic acid and plated into ultra-low attachment 6 well plates on an orbital shaker at 95rpm and fed daily. On day 14 the medium was switched to Hepatocyte Culture Medium (HCM, Lonza) with 10 ng/mL hepatocyte growth factor, 0.1uM dexamethasone and 20 ng/mL Oncostatin M. Cultures were fed daily until used for assays.

### Conditioned medium preparation

Cells were differentiated as described above. Subsequently they were exposed to HPLM medium (Thermo Fisher Scientific A4899101) for 5 days, with every other day feeding. Afterwards, HPLM medium was conditioned for 48 hours before harvesting, filtering, and being frozen at -80 C.

For the different varieties of adipocyte conditioned medium treated with small molecules, adipocytes were treated with the compound during the five-day HPLM exposure. Compounds were removed 48 hours prior to conditioned medium harvesting. Dexamethasone (Cayman Chemicals,11015) was used at 1uM, H151 (Avantor, MSPP- INH-H151) was used at 15uM, and diABZI (InVivoGen, tlrl-diabzi-2) was used at 1uM.

### Viral propagation and titration

West Nile virus (Nea Santa strain), Zika virus (Puerto Rico strain), Ilheus virus, and SARS-CoV-2 (isolate USA_WA1/2020) were obtained from BEI Resources. All viruses were propagated in Vero E6 cells (ATCC CRL-1586) cultured in Dulbecco’s modified Eagle’s medium (DMEM) supplemented with 2% fetal calf serum (FCS), penicillin (50 U/mL), and streptomycin (50 mg/mL). Viral titer was determined in Vero E6 cells by plaque assay. Work with SARS-CoV-2 was performed in the biosafety level 3 (BSL3) at the Ragon Institute (Cambridge, MA) following approved SOPs.

### Infection procedure

Target cells were incubated for 24 hours with either conditioned medium or unconditioned HPLM. At the time of infection, the cell culture medium was removed and replaced with inoculating virus diluted in a minimum volume of unconditioned HPLM. The cells were placed at 37°C for 1 hour at which point the inoculum was removed and replaced with conditioned or unconditioned HPLM. For neutralizing antibody experiments target cells were incubated with antibodies directed against either IFNα (InvivoGen Cat#hifna-mab1-02) or beta-galactosidase (InVivoGen Cat#bgal-mab1) for 30 minutes prior to infection. Antibodies were diluted at 1:5000 directly in conditioned medium. Following infection, the inoculum was removed and replaced with conditioned medium with or without neutralizing antibodies.

### Transwell experiments

Adipocytes were differentiated in 12-well plates. Approximately 1e5 differentiated hPSC-derived pericytes, macrophages, or dendritic cells were plated on the semipermeable membrane in the Transwell insert of Corning #3460 Transwell plates. 24 hours after plating cell culture medium was changed to HPLM and transwell inserts were placed in the 12-well plates containing adipocytes. 24 hours after the initiation of co-culture virus was added to the Transwell insert using the infection procedure described above. Medium was then collected from the top Transwell chamber at 48 hours post-infection.

### Immunofluorescence

Cells were fixed with 4% paraformaldehyde for 15 minutes at room temperature and washed three times with room temperature PBS. Fixed cells were permeabilized for 15 minutes using 0.1% TX-100. Cells were then blocked with 3% BSA in PBS for 1 hour at room temperature. Cells were incubated with primary antibodies (see Table 1) diluted in antibody buffer (0.01%TX-100 and 1%BSA in PBS) overnight at 4°C. Cells were washed with 0.05% Tween in PBS and incubated for 1 hour at room temperature with secondary antibodies (see Table 1) diluted in antibody buffer. DAPI was added at 1:10,000 with secondary antibodies.

### Cytokine Analysis

Conditioned medium was isolated from two independent differentiations of adipocytes, or adipocyte progenitors and sent to Eve technologies for cytokine analysis. Statistics were performed using GraphPad prism.

### Bulk RNA-sequencing

RNA was extracted from cells using the RNeasy Plus Mini kit (QIAGEN) following the manufacturer’s protocol. Libraries were prepared using the YourSeq (FT & 3’DGE) Strand-Specific mRNA Library Prep (Active Motif 23003) per manufacturer’ instructions following the FT protocol. All samples were sequenced on a NovaSeq 6000 sequencer.

### Analysis of gene expression

Paired-end reads (51x 51 bp) were mapped to the human genome (GRCh38) reference using the STAR aligner (v. 2.7.1a)^43^. Counts for human genes (ENSEMBL release 93 annotations), were tabulated using featureCounts^44^ with settings appropriate for strand-specific libraries. Differential expression of mRNAs was assessed for pairwise contrasts between conditions using estimated fold-changes and the Wald statistic in DESeq2 (v. 1.36.0)^45^. Per-gene dispersions were estimated using all conditions, as within-condition variance was observed to be relatively uniform across conditions in exploratory analyses. Unless otherwise noted, empirical Bayes shrinkage^46^ was applied to fold-change estimates. Gene-set enrichment analysis was carried out between pairs of conditions using GSEA (v. 4.1.0)^39^ with counts that were normalized by DESeq2 and 1000 gene-set permutations were used to estimate p-values. Gene-sets that were tested for enrichment came from the Hallmark collection^38^ at the MSigDB.

